# Expression of WIPI2B counteracts age-related decline in autophagosome biogenesis in neurons

**DOI:** 10.1101/325449

**Authors:** Andrea KH Stavoe, Erika LF Holzbaur

## Abstract

Autophagy defects have been implicated in multiple late-onset neurodegenerative diseases. Since aging is the most common risk factor in neurodegeneration, we asked how autophagy is modulated in aging neurons. We compared the dynamics of autophagosome biogenesis in neurons from young adult and aged mice, identifying a significant decrease in biogenesis during aging. Autophagosome assembly kinetics are disrupted, with frequent production of stalled isolation membranes in neurons from aged mice; these precursors failed to resolve into LC3-positive autophagosomes. We did not detect alterations in the initial induction/nucleation steps of autophagosome formation. However, we found that the transmembrane protein Atg9 remained aberrantly associated with stalled isolation membranes, suggesting a specific disruption in the WIPI-dependent retrieval of Atg9. Depletion of WIPI2 from young neurons was sufficient to induce a similar deficit. Further, exogenous expression of WIPI2 in neurons from aged mice was sufficient to restore autophagosome biogenesis to the rates seen in neurons from young adult mice, suggesting a novel therapeutic target for age-associated neurodegeneration.

## INTRODUCTION

Aging is a complex process that often impairs physiological and tissue function. Further, age is the most relevant risk factor for many prominent diseases and disorders, including cancers and neurodegenerative diseases (Niccoli and Partridge, 2012). Macroautophagy (hereafter referred to as autophagy) is an evolutionarily conserved, cytoprotective degradative process in which a double membrane engulfs intracellular cargo for breakdown and recycling (Cuervo et al., 2005; Rubinsztein et al., 2011). The autophagy pathway has been directly implicated in aging in model organisms (Cuervo, 2008; Rubinsztein et al., 2011). For example, *C. elegans* harboring autophagy mutations age faster and have shorter lifespans, whereas long-lived mutants display higher levels of autophagy (Hars et al., 2007; Kenyon et al., 1993; Meléndez et al., 2003; Tóth et al., 2008).

Neurons are post-mitotic, terminally differentiated cells which must maintain function in distal compartments throughout the lifetime of a human. These maintenance mechanisms may wane as a person ages, potentially contributing to neuronal dysfunction and death. Accordingly, misregulation of autophagy has been associated with several neurodegenerative diseases, including Alzheimer’s disease, Parkinson’s disease, Huntington’s disease, and amyotrophic lateral sclerosis (ALS) (Menzies et al., 2017; Nixon, 2013; Yamamoto and Yue, 2014). Furthermore, specifically disrupting autophagy in neurons results in neurodegeneration in animal models (Hara et al., 2006; Komatsu et al., 2006; Zhao et al., 2013). Despite the implication of this pathway in neurodegenerative disease, autophagy is best understood for its roles in maintaining cellular homeostasis in yeast and mammalian cells in response to stress (Abada and Elazar, 2014; Hale et al., 2013; Mariño et al., 2011; Reggiori and Klionsky, 2013; Son et al., 2012; Wu et al., 2013; Zhang and Baehrecke, 2015). Much less is known about how autophagy is regulated in neurons, where autophagy functions at a basal level (Hara et al., 2006; Komatsu et al., 2007; Yang et al., 2013; Yue et al., 2009) that is not significantly upregulated by proteomic stress (Maday et al., 2012; Wong and Holzbaur, 2014) or nutrient deprivation (Maday and Holzbaur, 2016). Importantly, little is known about how this essential homeostatic process is affected by aging.

While young neurons appear to clear dysfunctional organelles and protein aggregates very efficiently (Boland et al., 2008), few studies have examined autophagy in aged neurons. Since age is the most relevant risk factor in neurodegenerative disease (Niccoli and Partridge, 2012), elucidating how autophagy changes in neurons with age is crucial to understanding neurodegenerative diseases. Here, we examine how autophagy is altered with age in primary neurons from mice. We find that the rates of autophagosome biogenesis decrease in neurons with age. This decrease is not due to a change in the kinetics of either initiation or nucleation of autophagosomes. Instead, in neurons from aged mice, we find that the majority of autophagosome biogenesis events stall during autophagosome formation, failing to recruit LC3B with normal kinetics. Depletion of WIPI2 in neurons from young adult mice was sufficient to decrease the rate of autophagosome biogenesis to that of aged mice, while overexpression of WIPI2B in neurons from aged mice was sufficient to return the rate of autophagosome biogenesis to that found in neurons from young adult mice. Thus, while the rate of autophagosome biogenesis decreases in aged neurons, this decrease can be rescued by the restoration of a single autophagy component.

## RESULTS

### Autophagosome Biogenesis at the Axon Terminal of Neurites Decreases with Age

Autophagosome biogenesis, conserved from yeast to humans, involves over 30 proteins that act in distinct protein complexes to engulf either bulk cytoplasm or specific cargo within a double-membrane (Figure 1A). A signature autophagy protein, LC3B, has been used to label autophagosomes, as it becomes lipidated during autophagosome biogenesis and subsequently tightly associated with the limiting membranes of the autophagosome (Klionsky et al., 2016; Mizushima et al., 2010). We have previously used GFP-LC3B to examine autophagosome biogenesis in the distal tips of dorsal root ganglion (DRG) neurons from transgenic mice (Maday and Holzbaur, 2014; Maday et al., 2012). Unlike hippocampal or cortical neurons, which are typically isolated from embryonic or early postnatal rodents, DRG neurons can be isolated from mice of any age and grow robustly in culture following dissection. Importantly, the spatial enrichment of autophagosome biogenesis in the distal neurite observed in culture (Maday and Holzbaur, 2014; Maday et al., 2012) has been confirmed in *C. elegans* and *Drosophila* neurons *in vivo* (Neisch et al., 2017; Stavoe et al., 2016). Since impaired autophagy has been implicated in the pathogenesis of neurodegenerative diseases and age is the most relevant risk factor for neurodegenerative disease, we examined how autophagosome biogenesis rates change with age.

**Figure 1.**
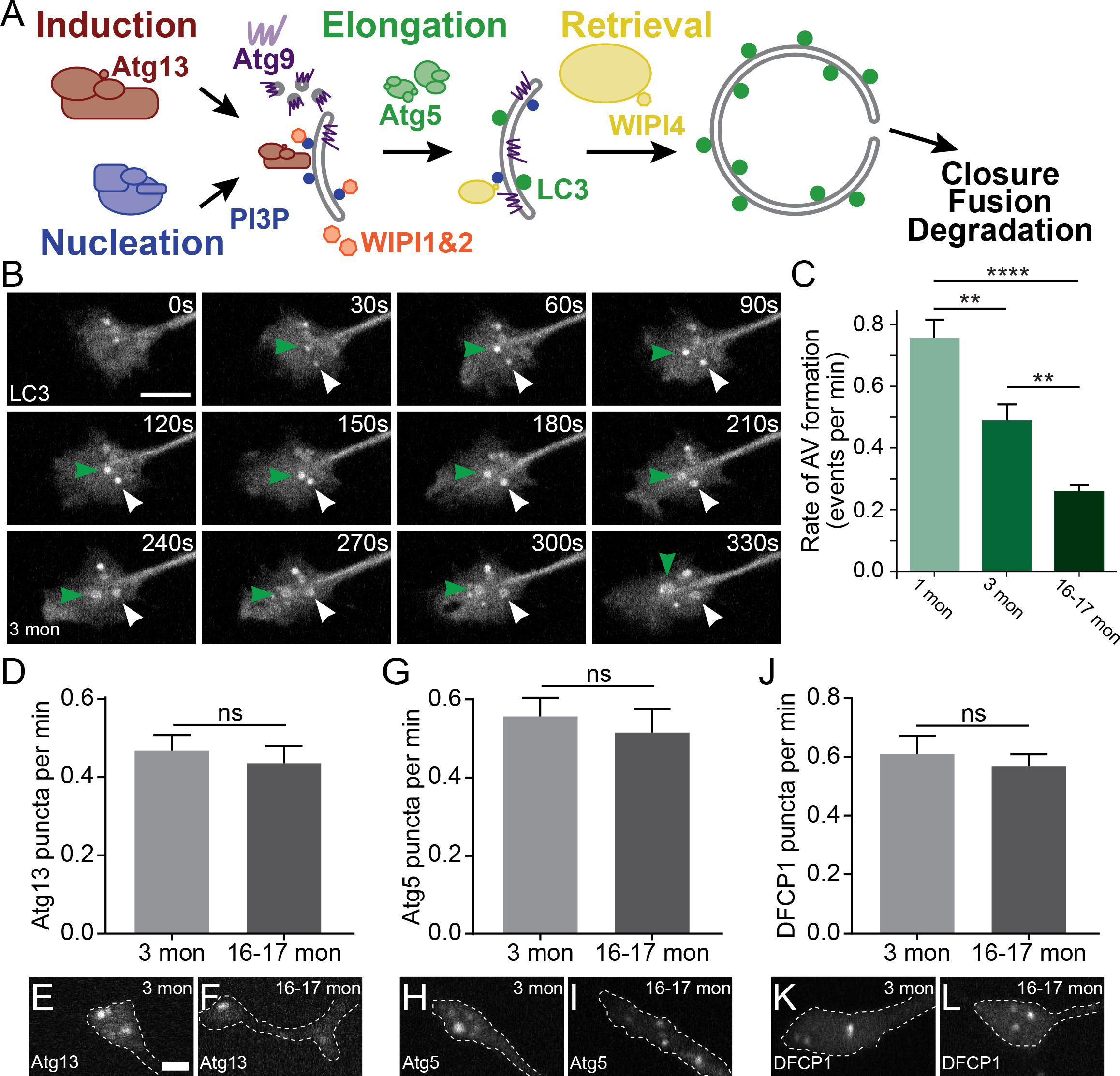
Autophagosome biogenesis decreases with age in neurons. (A) Schematic of autophagy pathway, focusing on the protein complexes involved in autophagosome biogenesis: induction (red), nucleation (blue), elongation (green), and retrieval (yellow). Atg9, a multi-pass transmembrane protein is in purple. The product of the nucleation complex, PI3P, is depicted as a blue dot, while LC3-II, the product of the elongation complex, is depicted as a green dot. WIPI1 and WIPI2, which bind to PI3P, are displayed in orange. (B) Time series of GFP-LC3 in the distal axon of DRG neurons from a young adult mouse. Green and white arrowheads each follow one autophagosome biogenesis event. Retrograde is to the right. Scale bar, 2 μm. (C) Quantification of the rate of autophagosome (AV) biogenesis in DRG neurons from young (1 mon, light green), young adult (3 mon, green), and aged (16-17 mon, dark green) mice (mean ± SEM; n≥ 54 neurons from three biological replicates). ** p < 0.01; ****p < 0.0001 by Kruskal-Wallis ANOVA test with Dunn’s multiple comparisons test. (D) Quantification of the rate of Atg13 puncta in DRG neurons from young adult (light gray) and aged (dark gray) mice (mean ± SEM; n≥ 28 neurons from three biological replicates). ns, not significant by Mann-Whitney test. (E-F) Representative micrographs of mCherry-Atg13 in the distal tip of DRG neurons from young adult (E) and aged (F) mice. (G) Quantification of the rate of Atg5 puncta in DRG neurons from young adult (light gray) and aged (dark gray) mice (mean ± SEM; n≥ 34 neurons from three biological replicates). ns, not significant by unpaired t test with Welch’s correction. (H-I) Representative micrographs of mCherry-Atg5 in the distal tip of DRG neurons from young adult (H) and aged (I) mice. (J) Quantification of the rate of DFCP1 puncta in DRG neurons from young adult (light gray) and aged (dark gray) mice (mean ± SEM; n≥ 56 neurons from three biological replicates). ns, not significant by unpaired t test with Welch’s correction. (K-L) Representative micrographs of Halo-DFCP1 in the distal tip of DRG neurons from young adult (K) and aged (L) mice. Scale bar in E, 2 μm, for E, F, H, I, K, L.

We assessed autophagosome biogenesis in primary DRG neurons dissected from mice at three different ages: postnatal day (P) 21-28 (one-month-old, young mice), P90-120 (three-month-old, young adult mice), or P480-540 (16-17-month-old, aged mice). We identified autophagosome biogenesis events as the formation of discrete GFP-LC3B puncta visible above the background cytoplasmic GFP-LC3 signal (Figure 1B). These puncta enlarge over approximately three minutes to form a 1μm autophagosome. We found that the rate of autophagosome biogenesis significantly decreases with age, with a rate of 0.76 events per minute (corresponding to 7.6 events within a ten minute video) in neurons from P21-28 mice, 0.49 events per minute in neurons from P90-120 mice, and 0.26 events per minute in neurons from P480-540 mice, corresponding to a 53% decrease in autophagosome biogenesis from young adult to aged neurons (Figure 1C).

Autophagosome biogenesis can be divided into five stages: initiation/induction, nucleation, elongation, retrieval, and membrane closure (Figure 1A). The initiation complex, including Atg13 and ULK1/Atg1, induces autophagosome biogenesis by phosphorylating other autophagy components (Feng et al., 2014; Kamada et al., 2000; Reggiori et al., 2004). The nucleation complex generates phosphatidylinositol 3-phosphate (PI3P) at the site of autophagosome biogenesis (Kihara et al., 2001; Obara et al., 2006). Subsequently, the elongation complex, composed of two conjugation complexes, is required to conjugate phosphatidylethanolamine (PE) to LC3 to yield LC3-II (Tanida et al., 2004). LC3-II is recruited to autophagosomes as the isolation membrane elongates during biogenesis and remains associated with autophagosomes until degradation of the internalized components. Atg9, a six-pass transmembrane protein, is thought to shuttle to the growing membrane with donor membrane (Koyama-Honda et al., 2013; Orsi et al., 2012; Sekito et al., 2009; Suzuki et al., 2015; Yamamoto et al., 2012; Young et al., 2006). The retrieval complex, including Atg2 and WIPI4, is thought to remove Atg9 from the vicinity of the growing autophagosome membrane (Reggiori et al., 2004; Wang et al., 2001). Finally, the limiting membrane closes and fuses with itself to generate the unique double-membrane organelle. The autophagosome then undergoes transport and subsequent fusion with lysosomes to degrade its contents (Xie and Klionsky, 2007).

The decrease in the rate of autophagosome biogenesis that we observed by monitoring GFP-LC3 could be the result of alterations to the initiation, nucleation or elongation complexes. To determine which stage of autophagosome biogenesis is affected by age, we compared the kinetics of each step of the pathway. We surveyed the role of the initiation complex by quantifying the appearance of mCherry-Atg13 puncta. We observed similar kinetics of Atg13 recruitment in neurons from young adult and aged mice (Figure 1D-1F). Thus, recruitment of the initiation complex does not change with age. Next, we examined elongation with an early autophagy marker, Atg5. The rate of puncta formation with the mCh-Atg5 marker also did not change with age (Figure 1G-1I), suggesting that recruitment of early elongation components is also not altered with age. Finally, to examine the effect of age on the nucleation complex, we used DFCP1, which binds to PI3P, the product of the autophagy nucleation complex (Figure 1A). Similar to other early autophagy steps, we did not detect a change in DFCP1 puncta formation with age (Figure 1J-1L), suggesting that the nucleation complex generates PI3P at the same rate during aging. Together, these data demonstrate that the decrease in the rate of autophagosome biogenesis with age is not due to an alteration in the kinetics of the early stages of autophagy.

### Pronounced stalling of autophagosome biogenesis is observed in aged neurons

We previously characterized the step-wise assembly kinetics of autophagosome biogenesis in neurons (Maday and Holzbaur, 2014). To determine whether altered step-wise kinetics during recruitment could contribute to the decrease in autophagosome biogenesis with age, we compared the assembly kinetics of Atg13, Atg5, and LC3 recruitment during autophagosome biogenesis. Surprisingly, we identified two populations of biogenesis events: “productive” events (Figure 2A, Movie S1), in which Atg13- or Atg5-positive puncta successfully recruit LC3, and “stalled” events (Figure 2B, Movie S2), in which Atg13- or Atg5-positive puncta fail to recruit LC3 during the imaging window (between five and ten minutes without successful recruitment). We note that these stalled autophagic vesicles (AVs) were still quite dynamic (Figure 2B, Movie S2). Using either Atg13 or Atg5 as a marker, we observed that neurons from young adult mice exhibited predominantly (~70%) productive biogenesis events, while neurons from aged mice displayed mostly (~70%) stalled events (Figure 2C-2D). Furthermore, we observed that both stalled and productive AVs could be observed within the same neuron and even in the same axonal tip (Figure S1, Movie S3). These data indicate that initial assembly of autophagy complexes is not altered with age, but rather that aging affects a later stage of autophagosome biogenesis, leading to an increase in stalled AVs in neurons from aged mice.

**Figure 2.**
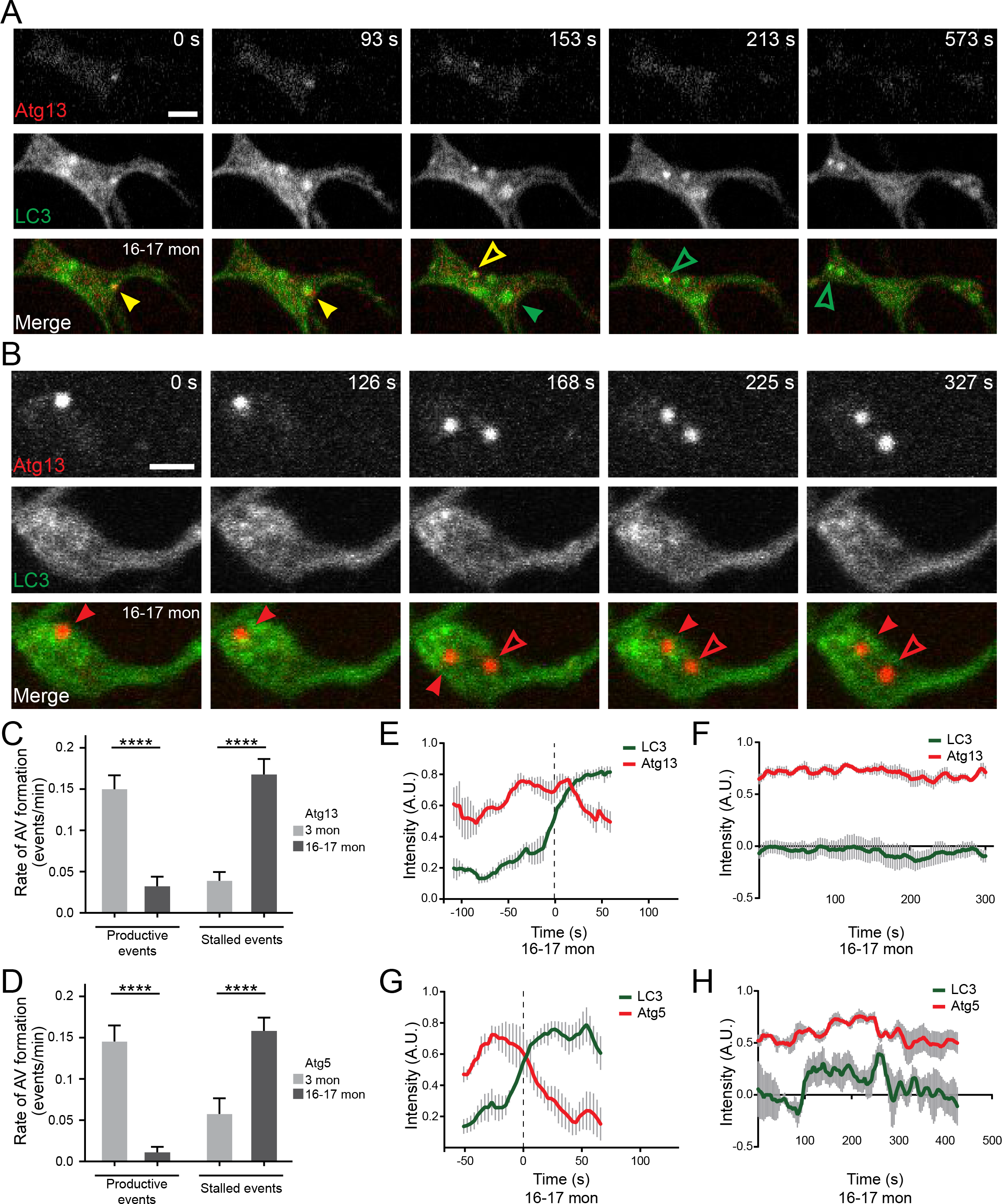
The ratio of stalled to productive autophagosome biogenesis events increases with age in neurons. (A) Time series of mCherry-Atg13 and GFP-LC3 in the distal neurite of DRGs from aged mice depicting a productive autophagosome biogenesis event. In the merge micrographs (bottom row), yellow arrowheads denote colocalization of Atg13 and LC3; green arrowheads denote a LC3-positive punctum from which Atg13 has dissociated; solid arrowheads track one punctum, hollow arrowheads follow a second punctum. Retrograde is to the right. Scale bar, 2 μm. (B) Time series of mCherry-Atg13 and GFP-LC3 in the distal neurite of DRG neurons from aged mice depicting a stalled autophagosome biogenesis event. In the merge micrographs (bottom row), red arrowheads denote a lack of colocalization between Atg13 and LC3 at Atg13-positive puncta that fail to recruit LC3; solid arrowheads track one punctum, hollow arrowheads follow a second punctum. Retrograde is to the right. Scale bar, 2 μm. (C-D) Quantification of the rate of AV formation (both productive and stalled events) in DRG neurons from young adult (light gray) and aged (dark gray) mice (mean ± SEM; n≥ 31 AVs from three biological replicates for each condition). AVs were identified with either Atg13 (C) or Atg5 (D). ****p < 0.0001 by Kruskal-Wallis ANOVA test with Dunn’s multiple comparisons test. (E-F) Mean intensity profiles of mCherry-Atg13 (red) and GFP-LC3 (green) for productive (E) and stalled (F) AVs (mean ± SEM; n≥ 5 biogenesis events from five neurons from three biological replicates). (G-H) Mean intensity profiles of mCherry-Atg5 (red) and GFP-LC3 (green) for productive (G) and stalled (H) AVs (mean ± SEM; n= 5 productive biogenesis events from five neurons or n=4 stalled biogenesis events from four neurons from three biological replicates). Vertical dashed line in (E and G) indicates the half-maximum of LC3 intensity, which was used to align the traces. See Figure S2A-S2H for intensity profiles of individual biogenesis events. See also Figures S1 and S2 and Movies S1-S3.

To assess the precise temporal relationship between Atg13, Atg5, and LC3 during autophagosome biogenesis, we quantified their fluorescence intensity over time. We first determined the assembly kinetics of LC3 compared to Atg13 (Figure 2E) or Atg5 (Figure 2G) in productive biogenesis events; we did not observe significant differences in the kinetics of productive events between neurons from young adult and aged mice. However, we did observe drastic differences in the kinetics of productive and stalled events in neurons from aged mice (Figure 2F and 2H, S2). A stalled event was defined as an Atg13- or Atg5- positive punctum (Atg13 or Atg5 remained 50% or more of max intensity in the video) that remained visible for five or more minutes without recruiting LC3 (LC3 signal at that punctum remained below 50% max LC3 intensity in the video). Taken together, these results suggest that aging does not impair the induction of autophagosome biogenesis, but instead impairs the subsequent recruitment of LC3 to the isolation membrane.

To further characterize the structure of the productive and stalled AVs, we used transmission electron microscopy to compare the ultrastructure of AVs at the axon terminals of neurons from young adult and aged mice. We observed stereotypical double-membrane structures containing cellular debris in the axonal tips of neurons from young adult mice (Figure 3A, 3C-3E, S3A-S3C). Distal tips of neurons from aged mice (Figure 3B) often contained more double-membrane-bound structures than distal tips of neurons from young adult mice (Figure 3A), and these occasionally contained cellular debris (Figure 3F). Interestingly, we observed that the predominant double-membrane-bound structures in aged neurons were multi-lammelar (Figure 3G, 3H, S3D-S3O), reminiscent of AVs observed in cortical biopsy specimens from patients with Alzheimer’s disease (Nixon et al., 2005).

**Figure 3.**
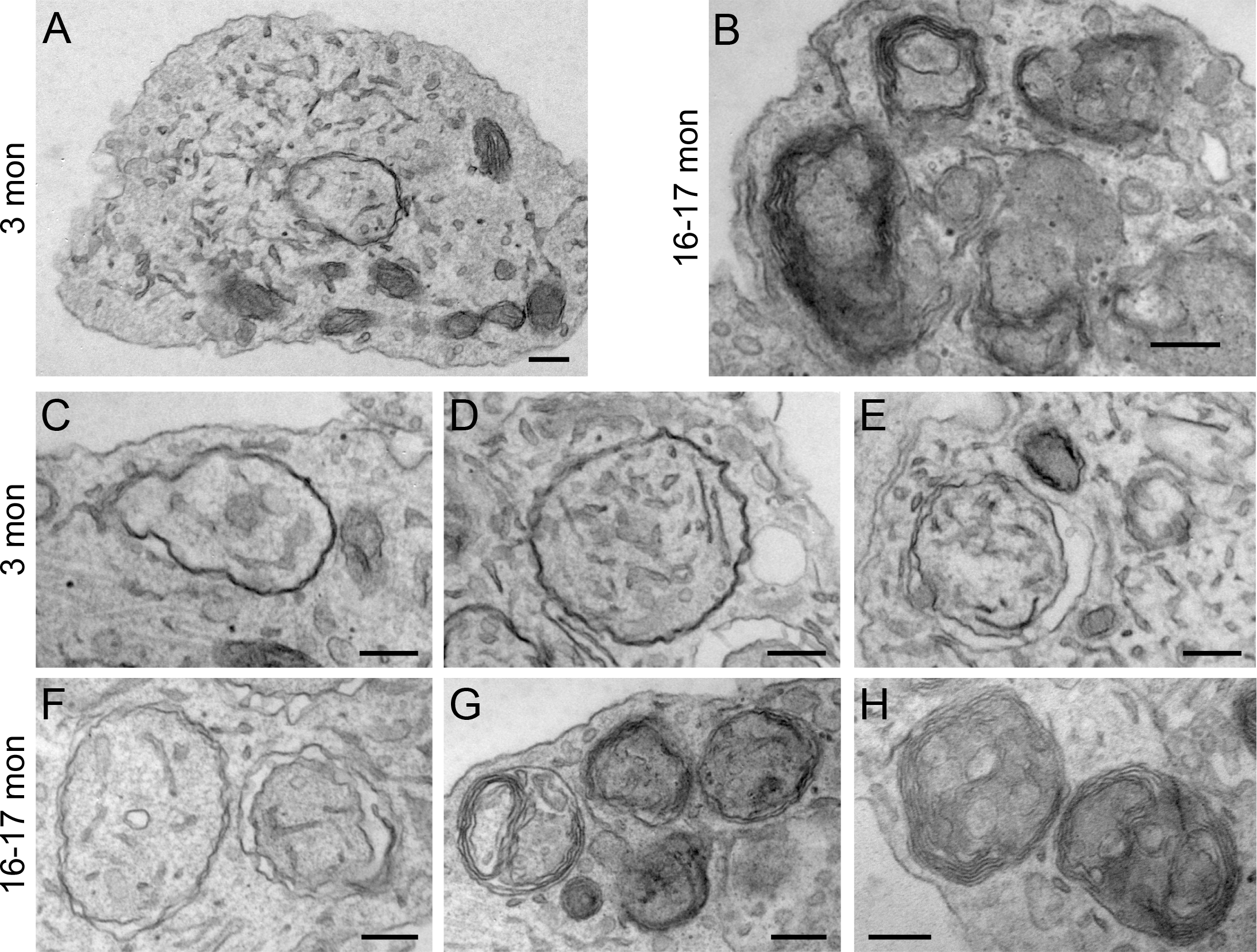
Autophagosomes display morphological differences via electron microscopy in neurons from young adult and aged mice. (A-B) Electron micrographs of DRG distal tips from young adult (A) or aged (B) mice. Scale bars, 200 nm. (C-E) Representative electron micrographs of autophagosomes in DRG distal tips from young adult mice. AVs are composed of a continuous double membrane enclosing engulfed cytoplasm. Scale bars, 200 nm. (F-H) Representative electron micrographs of autophagosomes in DRG distal tips from aged mice. AVs contain multiple, ruffled double membranes (G, H). Scale bars, 200 nm. See also Figure S3.

### Transmembrane Autophagy Protein Atg9 is Associated with a Majority of Stalled Biogenesis Events

Atg9 is the only multi-pass transmembrane protein in the core autophagy machinery (Feng et al., 2014; Lang et al., 2000; Noda et al., 2000; Young et al., 2006). While Atg9 is necessary for autophagy, its mechanistic role is unclear. It is thought that Atg9 transits to the growing isolation membrane with donor membranes (Sekito et al., 2009; Suzuki et al., 2015; Yamamoto et al., 2012; Young et al., 2006). Normally, Atg9 does not accumulate, but is only transiently recruited to the developing autophagosome (Koyama-Honda et al., 2013; Orsi et al., 2012). We asked whether Atg9 accumulates during stalled biogenesis events. We co-expressed SNAP-Atg9 with either mCh-Atg13 or mCh-Atg5 in neurons from young adult and aged mice. We observed that Atg9 colocalized with Atg13 or Atg5 at stalled AVs (Figure 4A, 4C, Movie S4). We scored for the presence of Atg9 at stalled AVs and determined that 60% of stalled events had associated Atg9. Of note, we also found that Atg9 remained associated with the relatively rare stalled AVs observed in neurons from young adult mice (Figure 4E). In contrast, productive AVs did not accumulate Atg9 in neurons from either young adult or aged mice (Figure 4B, 4D); only one out of 65 productive AVs was marked by Atg9. These data suggest that Atg9 may only transiently associate with productive biogenesis events, whereas the majority of stalled AVs aberrantly accumulate or retain Atg9.

**Figure 4.**
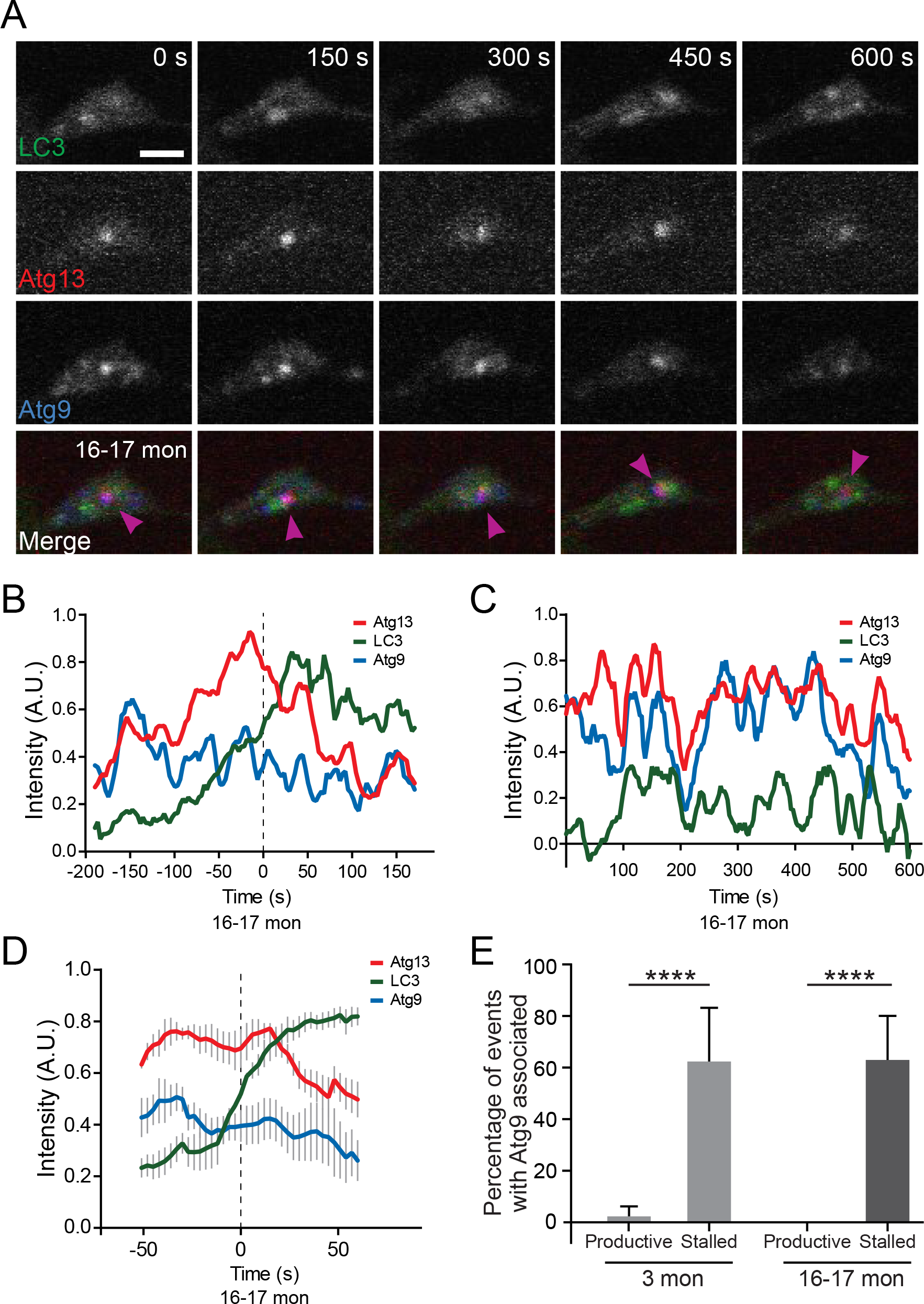
Atg9, a multi-pass transmembrane protein, associates with stalled autophagosome events. (A) Time series of mCherry-Atg13, SNAP-Atg9, and GFP-LC3 in the distal neurite of a DRG neuron from an aged mouse depicting a stalled AV. In the merge micrographs (bottom row), magenta arrowheads indicate colocalization between mCherry-Atg13 and SNAP-Atg9 without GFP-LC3. Retrograde is to the right. Scale bar, 2 μm. (B-C) Intensity profiles of mCherry-Atg13 (red), SNAP-Atg9 (blue), and GFP-LC3 (green) for a productive (B) and a stalled (C) AV in DRG distal tips from aged mice. (D) Mean intensity profiles of mCherry-Atg13 (red), SNAP-Atg9 (blue), and GFP-LC3 (green) for productive AVs (mean ± SEM; n= 5 biogenesis events from five neurons from three biological replicates). Vertical dashed lines in (B, D) indicate the halfmaximum of LC3 intensity, which was used to align the traces. (E) Quantification of the percentage of AVs that have Atg9 associated in the distal neurites of DRGs from young adult (light gray) and aged (dark gray) mice (n ≥ 17 for each age group). ****p < 0.0001 by two-tailed Fisher’s exact test. See also Movie S4.

### Knockdown of WIPI2 in neurons from young adult mice decreases the rate of autophagosome biogenesis toward that of neurons from aged mice

PROPPINs (β-propellers that bind phosphoinositides) are essential autophagy proteins that bind phosphotidyl-inositol-3-phosphate (PI3P) and are conserved from yeast to humans (Behrends et al., 2010; Michell et al., 2006; Polson et al., 2010; Proikas-Cezanne et al., 2004). In yeast, there is a single PROPPIN implicated in nonselective autophagy, Atg18 (Obara et al., 2008; Reggiori et al., 2004), while in mammals four family members have been identified. The mammalian PROPPINs are members of the WD-repeat protein interacting with phosphoinositides family (WIPI1 through WIPI4) (Polson et al., 2010; Proikas-Cezanne et al., 2004). Each WIPI protein binds two PI3P molecules via a highly conserved FRRG motif in the fifth and sixth β-propellers (Baskaran et al., 2012; Dove et al., 2004; Krick et al., 2006, 2012). WIPI2 and WIPI1 are highly homologous and considered to link PI3P production by the autophagy nucleation complex to LC3 conjugation to the isolation membrane (Dooley et al., 2014; Lamb et al., 2013; Polson et al., 2010). In addition, WIPI4 orthologs have been implicated in the removal of Atg9 from the isolation membrane, a process that is required for productive autophagosome biogenesis (Lu et al., 2011). Thus, we hypothesized that alterations in WIPIs may result in lower levels of productive autophagosome biogenesis.

We first tested whether siRNA depletion of WIPI2 levels in primary neuron culture would impair autophagosome biogenesis. We confirmed WIPI2 siRNA efficacy by western blot on cultured neurons from young adult mice treated with either control or WIPI2 siRNA (59% knockdown, Figure 5D). We observed that WIPI2 knockdown did not alter AV initiation, as marked by Atg5 (Figure 5B), but did significantly attenuate autophagosome biogenesis, indicated by LC3B recruitment. Rates of autophagosome formation fell from 0.39 to 0.09 AVs per minute (Figure 5C). We reintroduced siRNA-resistant human SNAP-WIPI2B, which did not alter the levels of autophagosome initiation, as marked by Atg5 (Figure 5B). However, rescue of WIPI2B levels restored the rate of autophagosome biogenesis to 0.41 AVs per minute, similar to levels seen in control neurons (Figure 5C). Thus, depleting WIPI2 in neurons from young adult mice reduces AV biogenesis, reminiscent of the age-dependent decrease in autophagosome biogenesis.

**Figure 5.**
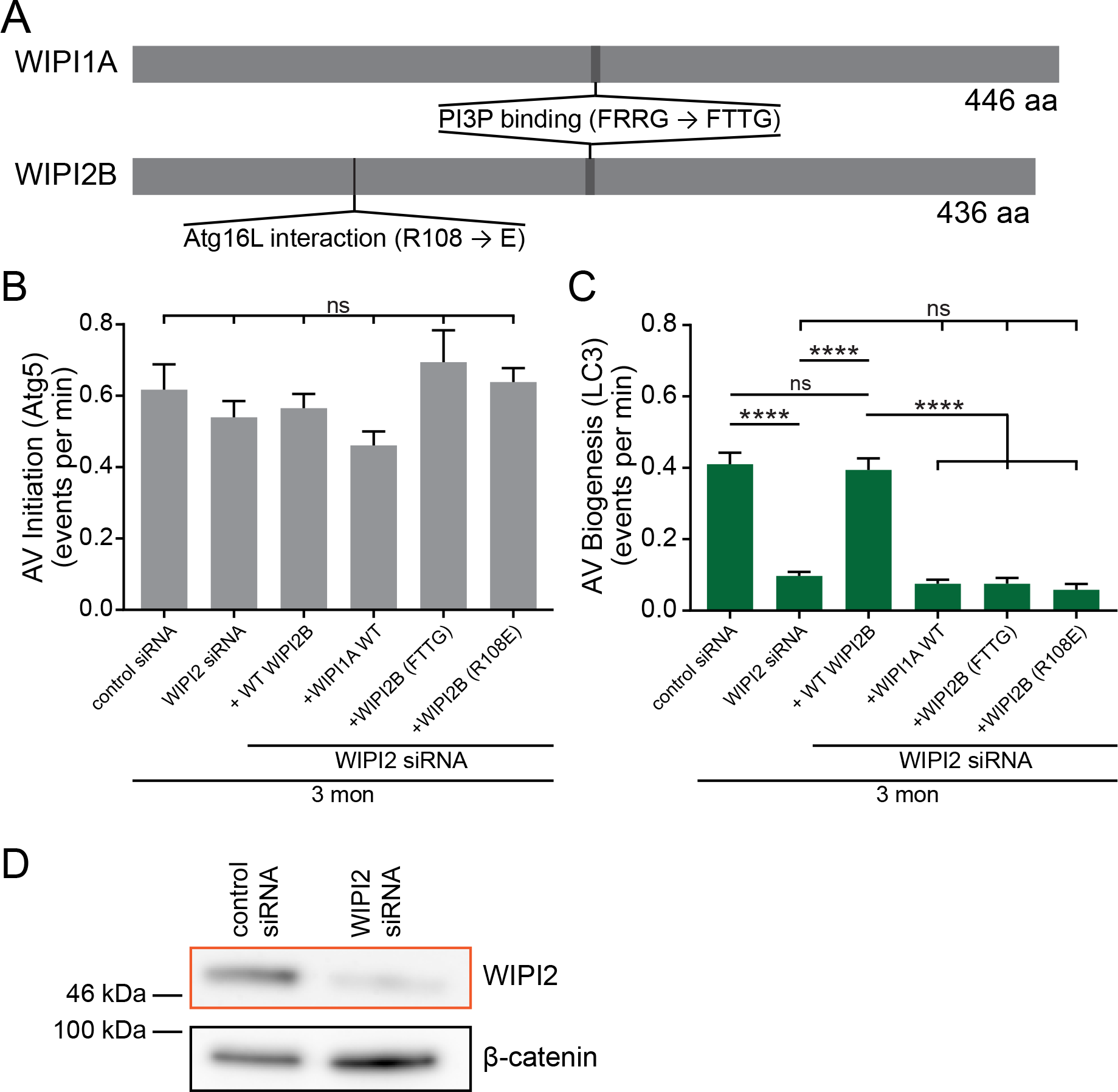
Knockdown of WIPI2 in neurons from young adult mice decreases autophagosome biogenesis. (A) Schematic of WIPI1A and WIPI2B proteins, depicting PI3P-interaction domains (FRRG) and Atg16L1-binding in WIPI2B (R108). Point mutations to disrupt these interactions are indicated in the relevant domains (FRRG → FTTG or R108 → E). (B) Quantification of the rate of AV initiation (mCh-Atg5 puncta) in DRG neurons from young adult with control or WIPI2 siRNA or WIPI2 siRNA with an RNAi-resistant WIPI2B construct (mean ± SEM; n≥ 15 neurons from three biological replicates for each siRNA condition). ns (not significant) p > 0.05 (Kruskal-Wallis ANOVA test with Dunn’s multiple comparisons test) (C) Quantification of the rate of AV biogenesis (GFP-LC3B puncta) in DRG neurons from young adult with control or WIPI2 siRNA or WIPI2 siRNA with an RNAi-resistant WIPI2B construct (mean ± SEM; n≥ 15 neurons from three biological replicates for each siRNA condition). ns (not significant) p > 0.05; ****p < 0.0001 by Kruskal-Wallis ANOVA test with Dunn’s multiple comparisons test. (D) Immunoblot of DRG lysates from young adult mice treated with indicated siRNA, collected after 2 days *in vitro*. β-catenin was used as a loading control. Equal total protein was loaded in each lane.

### Overexpression of WIPI2B in Neurons from Aged Mice Restores the Rate of Autophagosome Biogenesis to Young Adult Levels

Given that WIPI2 depletion impaired rates of autophagosome biogenesis in neurons from young adult mice, we hypothesized that overexpression of WIPI2 in neurons from aged mice would increase the rate of autophagosome biogenesis. In human, WIPI2 has five splice variants, but only two of these, including WIPI2B, are recruited to isolation membranes upon starvation in non-neuronal cells (Dooley et al., 2014; Mauthe et al., 2011; Polson et al., 2010). We found that Halo-WIPI2B colocalized with the early autophagosome marker mCh-Atg5 in neurons from aged mice (Figure 6A, Movie S5). While overexpression of WIPI2B in neurons from aged mice did not affect AV initiation (Figure 6B), ectopic expression of WIPI2B did increase rates of autophagosome biogenesis from 0.21 AVs per minute to 0.47 AVs per minute, a rate similar to that of neurons from young adult mice (Figure 6C). These more frequent productive biogenesis events in aged neurons exhibited similar autophagosome assembly kinetics as those in young adult neurons (Figure 6D). Consistent with its accumulation on stalled AVs, we observed that Atg9 did not accumulate at productive AVs in neurons from aged mice ectopically expressing WIPI2B (data not shown). These data suggest that overexpression of WIPI2B in neurons from aged mice restores autophagosome biogenesis and assembly kinetics to levels observed in neurons from young adult mice.

**Figure 6.**
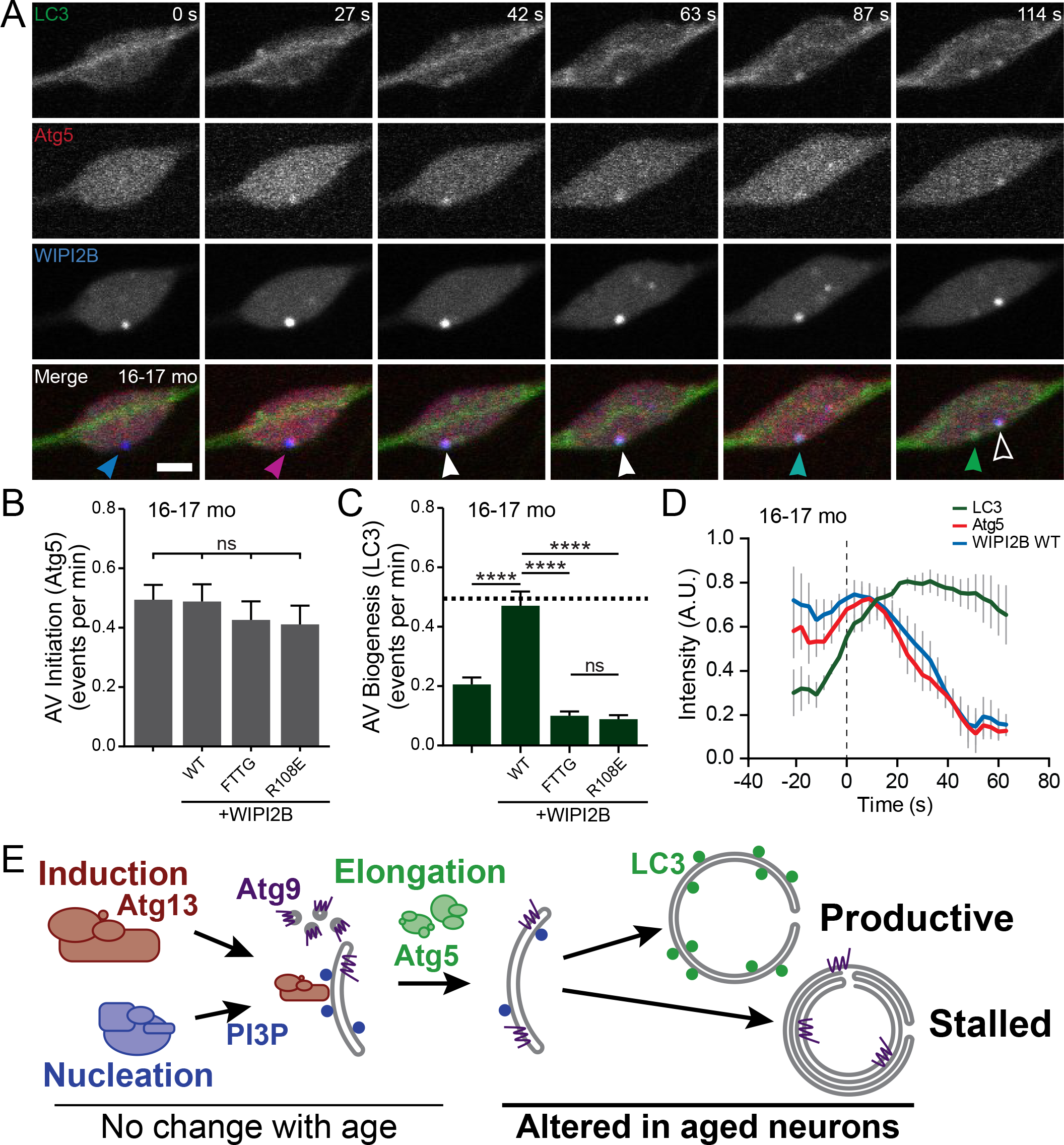
Overexpression of WIPI1 or WIPI2 in neurons from aged mice returns autophagosome biogenesis to levels observed in neurons from young adult mice. (A) Time series of GFP-LC3, mCherry-Atg5, and Halo-WIPI2B in the distal neurite of a DRG neuron from an aged mouse depicting a productive autophagosome biogenesis event. In the merge micrographs (bottom row), arrowheads indicate colocalization state on the isolation membrane; solid arrowhead follows one punctum, while outlined arrowhead indicates a different punctum. Retrograde is to the right. Scale bar, 2 μm. (B) Quantification of the rate of AV initiation (mCh-Atg5 puncta) in DRG neurons from aged (dark gray) with or without overexpression of WIPI2B (mean ± SEM; n≥ 15 neurons from three biological replicates for each condition). ns (not significant) p > 0.05 by Kruskal-Wallis ANOVA test with Dunn’s multiple comparisons test. (C) Quantification of the rate of AV biogenesis (marked with GFP-LC3B) in DRG neurons from aged mice with or without overexpression of indicated WIPI2B construct (mean ± SEM; n≥ 17 neurons from three biological replicates for each condition). **** p < 0.0001; ns (not significant) p > 0.05 by Kruskal-Wallis ANOVA test with Dunn’s multiple comparisons test. Horizontal dashed line indicates rate of AV biogenesis in neurons from young adult mice. (D) Mean intensity profiles of mCherry-Atg5 (red), Halo-WIPI2B (blue), and GFP-LC3 (green) for productive AVs (mean ± SEM; n= 6 biogenesis events from five neurons from three biological replicates). Vertical dashed line indicates the half-maximum of LC3 intensity, which was used to align the traces. See Figure S5A-S5C for intensity profiles of individual biogenesis events. (F) Model of how the autophagosome biogenesis pathway changes in neurons with aging. See also Movie S5.

### WIPI2B function in autophagosome biogenesis requires PI3P and Atg16L1 binding domains

Human WIPI2 and WIPI1 proteins are 80% identical, with further conservation in the lipid-binding motif (Polson et al., 2010) and in six out of seven WD40 repeats (Proikas-Cezanne et al., 2004). Due to the high homology between WIPI2 and WIPI1 (Figure 5A), we asked whether WIPI1A could substitute for WIPI2 upon WIPI2 knockdown in neurons from young adult mice. We found that expressing wild type WIPI1A upon WIPI2 depletion in neurons from young adult mice did not alter autophagy initiation (Figure 5B) and failed to rescue autophagosome biogenesis (Figure 5C). These data suggest that both WIPI2 and WIPI1 are required for autophagosome biogenesis and do not perform completely redundant functions.

Since WIPI2 and WIPI1 appear to have nonredundant functions, we asked how these highly homologous proteins differed. WIPIs have no known catalytic activity, but have distinct protein and lipid interaction domains. We were interested to determine how these interaction domains were required for WIPI function in autophagy. We first tested whether PI3P binding was required for WIPI2B function. As PROPPINs, WIPI2 and WIPI1 bind PI3P via a conserved FRRG motif within the fifth and sixth β-propellers (Baskaran et al., 2012; Dove et al., 2004; Gaugel et al., 2012; Jeffries et al., 2004; Krick et al., 2006; Proikas-Cezanne et al., 2004, 2007; Watanabe et al., 2012). This interaction can be abolished by mutating the positively charged arginine residues in the motif to uncharged threonine residues (FTTG) (Figure 7A) (Dooley et al., 2014).

We observed that autophagy initiation was not significantly altered upon ectopic expression of WIPI2B(FTTG) in neurons from young adult mice depleted of endogenous WIPI2 (Figure 5B). Similarly, expressing WIPI2B(FTTG) in neurons from aged mice did not affect AV initiation (Figure 6B). As we hypothesized, ectopic expression of siRNA-resistant WIPI2B(FTTG) in neurons from young adult mice treated with WIPI2 siRNA failed to rescue AV biogenesis (Figure 5C), which suggests that WIPI2B needs to bind to PI3P to promote autophagosome biogenesis. Additionally, overexpression of WIPI2B(FTTG) did not increase the basal rate of autophagosome biogenesis in neurons from aged mice (Figure 6C), further indicating that PI3P binding is required for WIPI2B function in autophagosome biogenesis. These results demonstrate that PI3P binding is required for WIPI2B function in autophagosome biogenesis.

Next, we focused on the Atg16L1-interaction domain of WIPI2B. In addition to PI3P binding, WIPI2B interacts with Atg16L1, an essential component of the LC3 conjugation complex. This function appears unique to WIPI2B, as WIPI1 cannot bind Atg16L1, likely due to the binding site being masked in WIPI1. This interaction between WIPI2B and Atg16L1 can be abrogated by switching a positively charged arginine to a negatively charged glutamate (R108E) in WIPI2B (Dooley et al., 2014).

We used a WIPI2B construct harboring a point mutation to eliminate WIPI2B binding to Atg16L1 (R108E) (Figure 5A) to determine whether binding to Atg16L1 is required for WIPI2B rescue of decreased autophagosome biogenesis. We did not observe changes in autophagy initiation with expression of the WIPI2B(R108E) construct in either WIPI2-depleted neurons from young adult mice (Figure 5B) or in neurons aged mice (Figure 6B). Interestingly, expression of WIPI2B(R108E) failed to rescue autophagosome biogenesis after WIPI2 depletion in neurons from young adult mice (Figure 5C). Similarly, ectopic expression of WIPI2B(R108E) in neurons from aged mice did not restore autophagosome biogenesis (Figure 6C), indicating that WIPI2B needs to interact with Atg16L1 to rescue decreased autophagosome biogenesis. These results confirm that both PI3P and Atg16L1 binding are necessary for WIPI2B function in autophagosome biogenesis. More importantly, our data indicate that the rate of autophagosome biogenesis decreases significantly over lifespan, but can be restored to rates seen in neurons from young adult mice by ectopic expression of WIPI2B.

## DISCUSSION

Here, we investigate the dynamics of autophagy during aging in primary neurons. We show that autophagosome biogenesis rates decrease in neurons with age. We further demonstrate that neither the initiation stage nor the nucleation stage of autophagy appears to be altered with age in mammalian neurons. Instead, we find that while some autophagosomes productively form in neurons from aged mice, the majority of autophagosome biogenesis events in aged neurons stall, failing to recruit LC3B with normal kinetics. Depleting WIPI2 in neurons from young adult mice was sufficient to decrease the rate of autophagosome biogenesis. Conversely, WIPI2B reconstitution in neurons from aged mice results in an increased rate of autophagosome biogenesis, restoring this rate to that found in neurons from young adult mice. Ultimately, we show that the rate of autophagosome biogenesis decreases in neurons during aging, but we can mitigate this decrease in cultured primary neurons from aged mice by overexpressing a single autophagy component, WIPI2B (Figure 6E).

Consistent with previous studies, our results indicate that WIPI1 and WIP2, two highly homologous PI3P-binding proteins, perform both redundant and unique functions (Dooley et al., 2014; Polson et al., 2010; Proikas-Cezanne et al., 2004). In neurons from aged mice, ectopic expression of WIPI2B is sufficient to return the rates of autophagosome biogenesis to young adult levels (Figure 6C). However, WIPI1A was not sufficient to rescue the rate of autophagosome biogenesis upon WIPI2 depletion in neurons from young adult mice (Figure 5C). These results suggest that WIPI1A is not fully capable of substituting for WIPI2B in autophagosome biogenesis. One important difference between WIPI1A and WIPI2B is that WIPI2B can bind to Atg16L1, a component of the elongation machinery, while WIPI1A cannot (Dooley et al., 2014), suggesting that this interaction between WIPI2B and the elongation complex is crucial to autophagosome biogenesis. It will be interesting to determine how these proteins change during lifespan, including protein levels and post-translational modifications.

Autophagosome biogenesis is only the initial step in this degradative pathway. Once autophagosome biogenesis is complete, the autophagosome matures by fusing with late endosomes and lysosomes, generating a proteolytically competent organelle. In neurons, this maturation occurs as the autophagosome is transported retrogradely to the cell soma, encountering lysosomes in transit (Fu et al., 2014; Hollenbeck, 1993; Lee et al., 2011a; Maday and Holzbaur, 2014, 2016; Maday et al., 2012; Wong and Holzbaur, 2014; Yue, 2007). Thus, in neurons, flux through the autophagy pathway is intimately linked with retrograde transport. The rate of autophagosome biogenesis will directly affect flux through the degradative pathway. Defects in autophagic flux (Lumkwana et al., 2017; Menzies et al., 2017; Nixon, 2013), AV transport (Tammineni et al., 2017), and lysosomal function (Gowrishankar et al., 2015; Lee et al., 2011a, 2011b) have been implicated in various neurodegenerative diseases. Furthermore, previous studies have identified mutations in organelle transport machinery that are linked to neurodegenerative disease (Maday et al., 2012; Millecamps and Julien, 2013). While we focused here on autophagosome biogenesis, it will be interesting to consider how autophagosome flux and transport through the axon may change with age in neurons. Additional age-based alterations in autophagosome flux and retrograde transport may compound the decline in rates of autophagosome biogenesis that we report here to result in age-dependent neurological deficits *in vivo*.

Our data indicate that in neurons from aged mice, the majority of the autophagosome biogenesis machinery assembles without recruiting LC3B (Figure 2); this failure to recruit LC3B correlates with stalled events. In mammals, there are multiple members of the LC3 and GABARAP subfamilies: LC3A, LC3B, LC3C, GABARAP, GABARAPL1/GEC1, and GABARAPL2/Gate-16 (Schaaf et al., 2016). Here, we examined the most studied family member, LC3B. Due to the high sequence similarity between LC3/GABARAP family members, it is thought that these proteins may perform similar, but not completely redundant functions in autophagy (Schaaf et al., 2016). It may be possible that other LC3/GABARAP family members act in autophagy at the distal tip in neurons from aged mice. However, the kinetics of the observed stalled AVs (Figure 2) suggest that none of the LC3/GABARAP family members are effectively recruited, preventing progression of the autophagy pathway. At an ultrastructural level, our data also indicate a change in AV morphology during aging, with a predominance of multiple-membrane-containing structures visible in neurons from aged mice (Figure 3). Together, these results suggest that LC3 recruitment may not be strictly required for autophagosome membrane generation and elongation. Instead, it is possible that LC3-II recruitment is required to regulate membrane generation to result in target engulfment by a solitary double-membrane structure. Without the recruitment of LC3B in neurons from aged mice, membrane elongation may continue unrestricted, generating the multiple-double-membrane structures observed by TEM in neurons from aged mice.

The autophagy pathway has been extensively studied in non-neuronal cell types, where autophagy can be induced by starvation or other cellular stress (Abada and Elazar, 2014; Hale et al., 2013; Mariño et al., 2011; Reggiori and Klionsky, 2013; Son et al., 2012; Wu et al., 2013; Zhang and Baehrecke, 2015). Conversely, *in vivo* and *in vitro* studies in neurons indicate that neuronal autophagy is not significantly induced by starvation (Fox et al., 2010; Maday and Holzbaur, 2016; Mizushima et al., 2004; Tsvetkov et al., 2010) or by proteotoxic stress (Maday et al., 2012; Wong and Holzbaur, 2014). These studies suggest that autophagosome biogenesis is regulated differentially in neuronal and non-neuronal cells. Our results indicate that induction of autophagosome biogenesis remains robust during aging in neurons, with early autophagosome markers accumulating with the same frequency in neurons from young adult and aged mice (Figure 1D-1L). Together, these data suggest that induction of autophagy in neurons is a relatively intrinsic feature that is independent of environmental factors. Therefore, it will be important to determine how neurons initiate autophagosome biogenesis with such precision in spatially disparate regions like axonal tips and during aging.

Misregulation of autophagy has been implicated in many neurodegenerative diseases and disorders. Aberrant accumulation of autophagosomes has been observed in axons in biopsies from Alzheimer’s disease patients (Nixon et al., 2005). Similarly, neuron-specific knockout of autophagy components results in neurodegeneration in mouse models (Komatsu et al., 2006, 2007). Furthermore, de novo mutations in WIPI4 are linked to neurodegeneration in adulthood (Haack et al., 2012; Saitsu et al., 2013). While ectopic induction of autophagy has met with some success in attenuating aggregated mutant huntingtin and Tau in neurodegeneration models (Ravikumar et al., 2002; Wang et al., 2009), it is still not clear whether protein accumulations are causally linked to neurodegeneration or simply correlated with aging (Rubinsztein et al., 2011). Our data indicate that autophagy pathway progression, not autophagy initiation (Figure 1D-1L), diminishes with age, suggesting that induction of autophagy may not be an effective target for treatment of neurodegenerative disease (Figure 6E). Rather, modulating later stages in autophagosome biogenesis may produce more successful therapies. Accordingly, future work will elucidate whether overexpressing WIPI2 in neurodegenerative models will yield a similar increase in rates of autophagosome biogenesis and result in tangible attenuation of neurodegeneration.

Here, we report that the rate of neuronal autophagosome biogenesis decreases during aging. Our results indicate that this is not due to a decline in autophagosome initiation, but to an increase in the number of stalled AVs that fail to recruit LC3B, and this defect can be attenuated by ectopically expressing WIPI2B. Together, these data suggest that an age-dependent disruption in a critical homeostatic pathway could be an over-arching commonality among different neurodegenerative disorders caused by many different factors. Further, our data demonstrate that the age-related decline in autophagosome biogenesis rates may be therapeutically reversible by overexpression of a single autophagy component, WIPI2, thus slowing the aging process in neurons.

## MATERIALS AND METHODS

### Reagents

GFP-LC3 transgenic mice (strain: B6.Cg-Tg(CAG-EGFP/LC3)53Nmi/NmiRbrc) were generated by N. Mizushima (Tokyo Medical and Dental University, Tokyo, Japan; (Mizushima et al., 2004)) and obtained from RIKEN BioResource Center in Japan. Constructs used include: mCherry-Atg5 (Addgene 13095), SNAP-Atg9 and Halo-Atg9 (subcloned from Addgene 60609), SNAP-WIPI1A (subcloned from Addgene 38272), and Halo-DFCP1 (subcloned from Addgene 38269). SNAP-WIPI2B and Halo-WIPI2B were subcloned from a generous gift from Sharon Tooze. SNAP-WIPI2B (FTTG), Halo-WIPI2B(FTTG), SNAP-WIPI2B(R108E), and Halo-WIPI2B(R108E) were generated via quick change and subcloned into original plasmids. The SNAP backbone was originally obtained from New England Biolabs (NEB), and the Halo backbone was originally obtained from Promega.

### Primary Neuron Culture

DRG neurons were isolated using the same protocol as (Perlson et al., 2009) and cultured in F-12 Ham’s media (Invitrogen) with 10% heat-inactivated fetal bovine serum, 100 U/mL penicillin, and 100 μg/mL streptomycin. For live-cell microscopy, DRGs were isolated from P21-28 (young), P90-120 (young adult), or P480-540 (aged) mice and plated on glass-bottomed dishes (MatTek Corporation) and maintained for 2 days at 37°C in a 5% CO_2_ incubator. Prior to plating, neurons were transfected with a maximum of 0.6 μg total plasmid DNA using a Nucleofector (Lonza) using the manufacturer’s instructions. Microscopy was performed in low fluorescence nutrient media (Hibernate A, BrainBits) supplemented with 2% B27 and 2 mM GlutaMAX. For nucleofected constructs that yielded Halo- or SNAP-tagged proteins, DRGs were incubated with 100nM of the appropriate Halo or SNAP ligand (SNAP-Cell^®^ 647-SiR, SNAP-Cell^®^ TMR-Star, and SNAP-Cell^®^ 430 from NEB; HaloTag^®^ TMR Ligand from Promega; silicon-rhodamine-Halo ligand from K. Johnsson, École Polytechnique Federale de Lausanne, Lausanne, Switzerland) for at least 30 min at 37°C in a 5% CO_2_ incubator. After incubation, DRGs were washed three times with complete F-12 media, with the final wash remaining on the DRGs for at least 15 min at 37°C in a 5% CO_2_ incubator.

Mice of either sex were euthanized within the indicated postnatal range. All animal protocols were approved by the Institutional Animal Care and Use Committee at the University of Pennsylvania.

### Live-Cell Imaging and Image Analysis

DRG microscopy was performed on a spinning-disk confocal (UltraVIEW VoX; PerkinElmer) microscope (Eclipse Ti; Nikon) with an Apochromat 100x, 1.49 NA oil immersion objective (Nikon) at 37°C in an environmental chamber. The Perfect Focus System was used to maintain Z position during time-lapse acquisition. Digital micrographs were acquired with an EM charge-coupled device camera (C9100; Hammamatsu Photonics) using Volocity software (PerkinElmer). Time-lapse videos were acquired for 10 min with a frame every 3 sec to capture autophagosome biogenesis. Multiple channels were acquired consecutively, with the green (488 nm) channel captured first, followed by red (561 nm), far-red (640 nm), and blue (405 nm). DRGs were selected for imaging based on morphological criteria and low expression of transfected constructs.

Time-lapse micrograph stacks were analyzed with FIJI (Schindelin et al., 2012). “Stalled” biogenesis events were defined as Atg13 or Atg5 puncta that remained visible for at least five minutes and never recruited LC3 within the 10 min video. “Productive” biogenesis events were defined as Atg13/Atg5 puncta that did recruit LC3 within in the ten minute video.

### Biochemistry

Lysates were prepared from brains or DRGs of non-transgenic mice. Brains were homogenized individually in RIPA buffer [50mM NaPO4, 150 mM NaCl, 1% Triton X-100, 0.5% deoxycholate, 0.1% SDS, and 1x complete protease inhibitor mixture (Roche)]. DRGs were homogenized in RIPA buffer with a 1.5mL pestle. Homogenized samples were lysed for 30 minutes on ice. For the siRNA controls, isolated DRGs were plated at 60,000 neurons per 35 mm dish as described above for 2 DIV. Neurons were washed with PBS (50mM NaPO4, 150 mM NaCl, pH 7.4) and then lysed as above. Samples were centrifuged at 17,000 × g at 4°C for 15 minutes. Resultant supernatants were analyzed by SDS-PAGE, transferred onto Immobilon P PVDF membrane, and visualized with enhanced chemiluminescent substrate (SuperSignal West Pico chemiluminescent substrate, ThermoFisher Scientific) on the G:Box and GeneSys digital imaging system (Syngene). Antibodies used include: a polyclonal antibody against LC3B (Abcam ab48394, 1:1000), a monoclonal antibody against WIPI2 (Abcam ab105459, 1:1000), a polyclonal antibody against Atg9A (Abcam ab105402, 1:250), a polyclonal antibody against WIPI1 (Thermo Fisher PA5-34973, 1:500), a monoclonal antibody against actin (Millipore MAB1501, 1:3000), and a monoclonal antibody against β-catenin (BD Biosciences 610154, 1:5000).

Bands from western blots were analyzed with the gel analyzer tool in FIJI (Schindelin et al., 2012). In each lane, the value of the protein of interest was divided by the value of the corresponding loading control (β-catenin or actin) to control for differences in sample loading. Ratios were then normalized to young adult averages and expressed as fold change of control.

### Electron Microscopy

Non-transgenic DRGs were isolated as above and plated as spot cultures on glass-bottomed dishes (MatTek Corporation) and maintained for 2 days at 37°C in a 5% CO_2_ incubator. DRGs were fixed with 2.5% glutaraldehyde, 2.0% paraformaldehyde in 0.1M sodium cacodylate buffer, pH 7.4, overnight at 4°C. Fixed DRGs were then transferred to the Electron Microscopy Resource Laboratory at the University of Pennsylvania, where all subsequent steps were performed. After subsequent buffer washes, the samples were post-fixed in 2.0% osmium tetroxide for 1 hour at room temperature and then washed again in buffer, followed by dH_2_O. After dehydration through a graded ethanol series, the tissue was infiltrated and embedded in EMbed-812 (Electron Microscopy Sciences, Fort Washington, PA). Thin sections were stained with lead citrate and examined with a JEOL 1010 electron microscope fitted with a Hamamatsu digital camera and AMT Advantage image capture software. Regions between cell body densities with maximum neurite invasion were chosen for imaging.

### Additional Methods

All image analysis was performed on raw data. Images were prepared in FIJI (Schindelin et al., 2012); contrast and brightness were adjusted equally to all images within a series. Figures were assembled in Adobe Illustrator. Prism (GraphPad) was used to plot graphs and perform statistical tests; for normally distributed data, one-way ANOVAs were performed with Tukey’s multiple comparison’s test; for nonparametric data, one-way ANOVAs were performed with Kruskal-Wallis test.

## AUTHOR CONTRIBUTIONS

A.K.H.S. and E.L.F.H. designed the experiments. A.K.H.S. performed and analyzed the experiments, and A.K.H.S. and E.L.F.H. prepared the manuscript.

## ACKNOWLEDGEMENTS

We gratefully thank Mariko Tokito and Karen Jahn for their technical assistance and the members of the Holzbaur lab for thoughtful discussions, encouragement, and comments on the manuscript. This work is supported by funding from NIH grant R37 NS060698 to E.L.F.H. A.K.H.S. is supported by NIH fellowship F32 NS100348-01.

